# Canalization of circuit assembly by δ-protocadherins in the zebrafish optic tectum

**DOI:** 10.1101/2025.01.29.635523

**Authors:** Sayantanee Biswas, Michelle R. Emond, Grace S. Philip, James D. Jontes

## Abstract

Neurons are precisely and reproducibly assembled into complex networks during development. How genes collaborate to guide this assembly remains an enduring mystery. In humans, large numbers of genes have been implicated in neurodevelopmental disorders that are characterized by variable and overlapping phenotypes. The complexity of the brain, the large number of genes involved and the heterogeneity of the disorders makes understanding the relationships between genes, development and neural function challenging. Waddington suggested the concept of canalization to describe the role of genes in shaping developmental trajectories that lead to precise outcomes^1^. Here, we show that members of the δ-protocadherin family of homophilic adhesion molecules, Protocadherin-19 and Protocadherin-17, contribute to developmental canalization of visual circuit assembly in the zebrafish. We provided oriented visual stimuli to zebrafish larvae and performed *in vivo* 2-photon calcium imaging in the optic tectum. The latent dynamics resulting from the population activity were confined to a conserved manifold. Among different wild type larvae, these dynamics were remarkably similar, allowing quantitative comparisons within and among genotypes. In both Protocadherin-19 and Protocadherin-17 mutants, the latent dynamics diverged from wild type. Importantly, these deviations could be averaged away, suggesting that the loss of these adhesion molecules leads to stochastic phenotypic variability and introduced disruptions of circuit organization that varied among individual mutants. These results provide a specific, quantitative example of canalization in the development of a vertebrate neural circuit, and suggest a framework for understanding the observed variability in complex brain disorders.

## Introduction

Despite their immense complexity, vertebrate brains are highly organized and exhibit a conserved, stereotyped architecture. Distinct brain regions comprise canonical circuits that carry out well-defined functions, which are conserved both within and between species^2,3^. The assembly of these conserved circuits is robust to genetic and environmental variation, as well as natural biological heterogeneity. This allows the brains of distinct individuals to perform the same tasks, even though they are not identical at the level of neurons and synapses, or in their particular developmental histories. The mechanisms that confer robust circuit development in the face of these multiple sources of variability are poorly understood^4,5^.

In the case of neurodevelopmental disorders, such as schizophrenia and autism, the relationship between conserved structure and conserved function breaks down. Characterized by profound disturbances in normal brain function, some evidence suggests that these disorders are “connectopathies,” arising from disruption of the underlying network topology^6–10^. While a large number of genes have been linked to developmental brain disorders, understanding the relationship of genotype to phenotype is challenging. In addition to natural biological variability, these disorders are plagued by variability in penetrance and expressivity^11,12^. Moreover, putatively distinct disorders have amorphous boundaries, overlapping with one another and exhibiting comorbidities^13^, as well as sharing genetic origins^14–16^. Collectively, these sources of variation mask the relationships between gene function, circuit organization and neural activity. These same issues make interpretation of studies in animal models similarly challenging.

Protocadherin-19 is a member of the δ-protocadherin family of homophilic cell adhesion molecules^17–19^. In humans, mutations in *PCDH19* are linked to schizophrenia and autism^20,21^, and cause a developmental epileptic encephalopathy^22,23^. To investigate how mutations in *pcdh19* or the related *pcdh17* affect neural development, we used visual stimulation and *in vivo* 2-photon calcium imaging in the optic tectum of the larval zebrafish. We show that the neuronal population responses to oriented sinusoidal gratings define neural dynamics that are strongly stereotyped among wild type larvae. In mutants, these dynamics are altered, reflecting stochastic deviations in circuit organization. Our results are consistent with a role for protocadherins in canalization of development, contributing to robustness of circuit assembly. This can provide a conceptual framework for understanding the variability associated with neurodevelopmental disorders.

## Results

Moving stimuli, such as moving lines, spots or gratings, elicit robust responses in the optic tectum of zebrafish^24–27^. To record the activity of tectal neurons, we generated a transgenic line that *pan*-neuronally expresses a nuclear localized jGCaMP8s (Histone H2B-GCaMP8s). These transgenic larvae were mounted in a custom imaging chamber **(Fig. 1a**), and we elicited visual responses by projecting a sequence of drifting sinusoidal gratings (12 directions rotated at 30° intervals) onto an adjacent translucent screen (**Fig. 1b,c**). Two-photon image stacks spanning the optic tectum (7 optical sections at 10 μm spacing) were collected at 1 Hz (**Fig. 1d**), allowing us to sample population activity. We used constrained non-negative matrix factorization (CNMF) with CaImAn^28^ to automatically segment images and extract fluorescence traces from visually-responsive neurons (**Fig. 1e,f**). For further analysis, the neural responses were averaged across three trials (**Fig. 1g**). Overall, we obtained 2506 visually responsive neurons (n=9).

**Figure 1.**
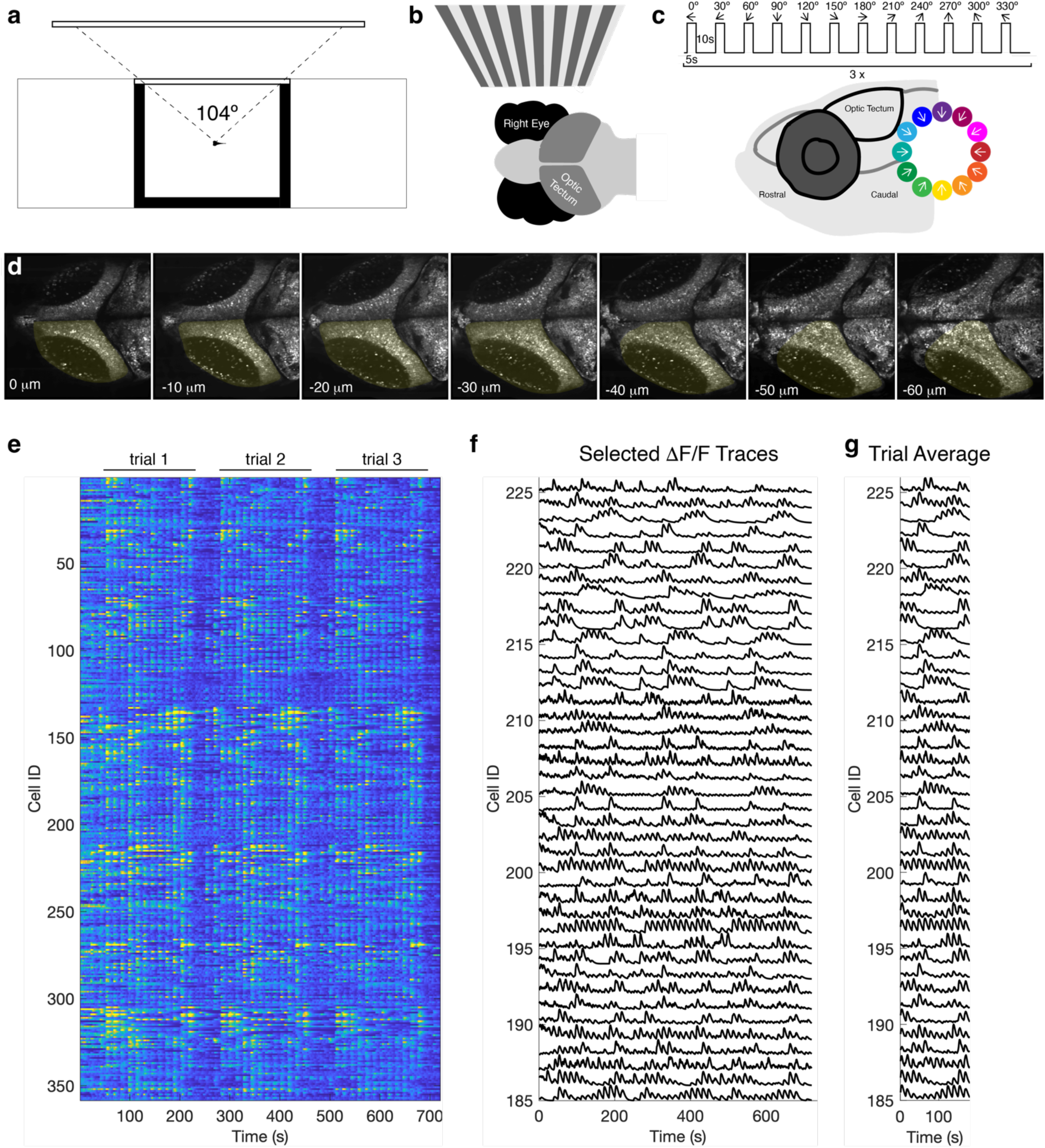
Calcium-imaging of neural responses to oriented sinusoidal gratings. **a.** Schematic of the experimental setup. A 6dpf larva is embedded in agarose in an imaging chamber, allowing 104° view of a projection screen. **b.** The visual stimuli are presented to the right eye and the neural responses are visualized in the contralateral (left) hemi-tectum. **c.** The stimulation protocol is a series of 5s on 10s off of a moving sinusoidal grating, 12 directions spaced at 30° intervals to encompass the full 360°. This is repeated three times. Below is a lateral view of the head of a zebrafish larva. The inset shows the directions of motion of sinusoidal gratings with reference to the rostro-caudal axis of the fish. The direction color-code is used throughout the paper. **d.** Image stacks consisting of 7 optical sections spaced at 10 μm were collected at 1s intervals. Shown are average intensity images of individual planes from a timelapse sequence collected in a 6 dpf transgenic larva, Tg(elav3l:HistoneH2B-GCaMP8s). The visually-responsive hemi-tectum is outlined in yellow. **e.** Example data showing all the fluorescence traces from visually-responsive neurons collected from one larva. **f.** ΔF/F traces selected from those shown in **e**. **g.** Responses were averaged across the three trials. Shown are the trial averages from the traces in **f**.

It is now widely understood that sensory processing and motor control are encoded by neuronal population dynamics, rather than the activities of individual neurons^29,30^. These latent dynamics consist of a small number of covariation patterns, or neural modes, which describe a neural manifold within a low-dimensional subspace^31–33^. To explore the structure of tectal population dynamics in response to visual stimulation, we performed linear dimensionality reduction with principal component analysis (PCA) (**Fig. 2a**). For *n* neurons, PCA reduces the data from an *n*-dimensional space to a more compact *k*-dimensional space. We found that the first five neural modes captured 89.9±1.8% (n=9; mean ± sem.) of the variance in our wild type data (**Fig. 2b**), and our subsequent analyses are confined to these five neural modes. Each axis represents a significant pattern of neuronal covariation that is activated to varying extent during the presentation of visual stimuli, and projection of the original neural data onto these axes describes the latent dynamics evoked by visual stimulation (**Fig. 2c**). To determine how consistent the neural responses are in individual larvae, the latent dynamics were obtained for individual trials, and the Pearson’s correlation (*r*) between individual trials was calculated along each dimension (**Fig. 2c,d**). The mean trial-to-trial correlation was greater than 0.9 across all 5 neural modes (**Fig. 2d**). Recent evidence suggests that the similarity of neural architecture among individuals gives rise to similar latent dynamics for a given stimulus or motor task^34^. In mammals, latent dynamics are estimated from a small subsample of neurons, which varies from individual to individual. In such cases, the latent dynamics from different animals need to be aligned using a procedure such as canonical correlation analysis^34,35^. Here, we image a relatively large proportion of visually-responsive neurons and sample among the same neuronal types across different fish, obviating the need for alignment. For each wild type individual, we determined the latent dynamics from trial averaged responses (**Fig. 2e**). To quantify the similarity of the dynamics among the pool of wild types, we generated the averaged response, then calculated Pearson’s correlation between this average and the dynamics of each individual (**Fig. 2e,f**). The mean correlation along each neural mode was nearly as high as for trial-to-trial correlations (**Fig. 2c,d**), indicating that the similarity of dynamics among individuals is comparable to the similarity of successive trials in the same individual. Thus, the latent dynamics elicited by visual stimulation is stereotyped across different zebrafish larvae, facilitating comparisons within and among experimental groups. While the latent dynamics among wild type larvae are strongly correlated, it is possible that the correlation could be improved by further alignment. To test this, we performed Canonical Correlation Analysis (CCA) on our wild type datasets, but found that alignment degraded the correlation of dynamics derived from wild type larvae (*not shown*). When plotted in three dimensions, the latent dynamics from individual larvae describe a rosette shaped trajectory, with each leaf representing the response to a direction of grating drift (**Fig. 2g,h**). The averaged dynamics yielded a smooth trajectory in three dimensions (**Fig. 2i**). Thus, the population response to our simple stimulus presentation, encodes the direction of motion on a neural manifold that is preserved across zebrafish larvae.

**Figure 2.**
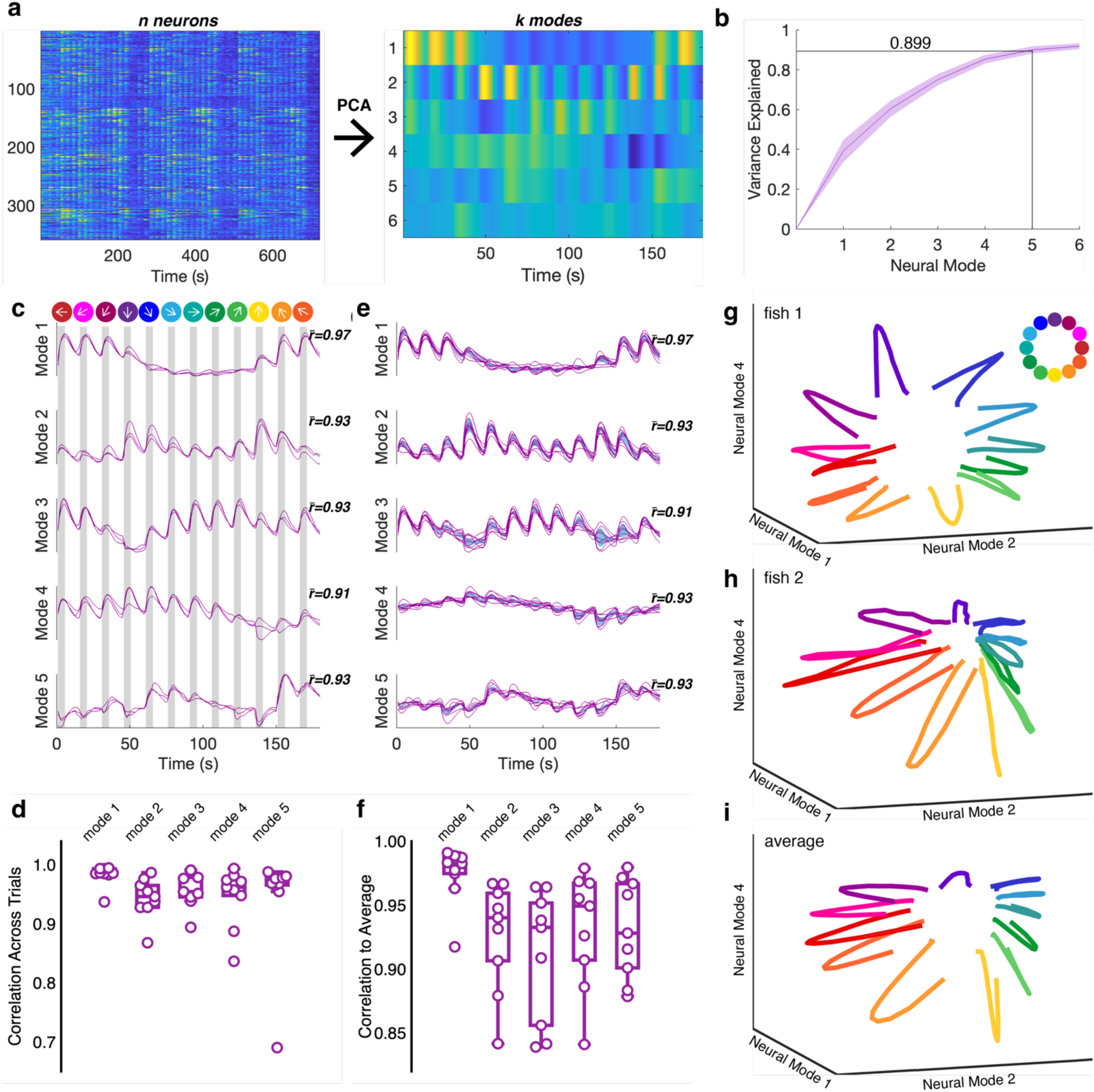
Latent dynamics in the optic tecta of zebrafish larvae in response to visual stimulation. **a.** Illustration of our approach to data analysis. Datasets such as the one shown on the left consisted of fluorescence traces from *n* neurons. These were trial averaged and the dimensionality of the trial averaged data was reduced using PCA. The new *k* dimensions are referred to as neural modes. **b.** The first 5 neural modes explain ∼90% of the variance in the data. The shaded area represents the standard deviation, n=9). **c.** Shown are the latent dynamics of an individual wild type larva for the first 5 neural modes. The latent dynamics were calculated independently for each of the three trials. The mean correlation of the dynamics among the individual trials to the trial average is shown as *r*’. The presentation of each stimulus is shown as a gray bar, and the direction of each stimulus is shown at the top. **d.** Graph summarizing the mean correlations across trials along each mode for the wild type larvae (0.98±0.006, mode 1; 0.94±0.012, mode 2; 0.96±0.01, mode 3; 0.95±0.017, mode 4; 0.94±0.032, mode 5 ;n=9) **e.** Latent dynamics determined from trial averaged data. Traces (purple) are the dynamics from individual larvae and the shaded area (blue) is the variance for the wild type group. The value, *r*’, represents the mean correlation of each individual neural mode to the mean dynamics averaged across the group (see **f**). **f.** Summary of the correlation between individual larvae and the group average along neural modes (0.97±0.008, mode 1; 0.93±0.014, mode 2; 0.91±0.017, mode 3; 0.93±0.015, mode 4; 0.93±0.013, mode 5; n=9). **g,h.** Three-dimensional representation of latent dynamics for two, individual larvae. Each color represents a direction of motion for the visual stimulus. The excursions of neural trajectories during stimulus presentation describes a roughly circular neural manifold. **i.** A three-dimensional representation of latent dynamics averaged across the wild type group (n=9).

To investigate the involvement of *pcdh19* in the assembly of visual circuitry we used two, distinct *pcdh19* mutant alleles. We previously introduced an indel lesion near the 5’ end of exon1 shortly after the sequence encoding the signal peptide^36,37^, resulting in a complete lack of functional Pcdh19 (*pcdh19^-^* ^10b^*^p^*). In addition, we have generated a new “promoterless” *pcdh19* allele, which lacks ∼4kb of genomic sequence, spanning the basal promoter, 5’UTR and the ATG start codon in exon 1 (*pcdh19^Δprom^*)(**Fig. S1a,b**). While both alleles completely eliminate the production of Pcdh19 protein (**Fig. S1c**), evidence suggests that indel mutations, like *pcdh19^-^*^10b^*^p^*, can elicit compensatory mechanisms arising from nonsense-mediated RNA decay^38,39^. Both of these mutant lines were crossed into our *Tg(elav3l:H2B-GCaMP8s)* line. As above, we performed calcium imaging on homozygous mutants during visual stimulation, and computed the latent dynamics for each mutant from the trial averages of visually responsive neurons (**Fig. 3a,b**). To determine whether the mutant groups differed from wild type in their visual responses, we correlated the trial-averaged dynamics from each individual mutant with the wild type group average along each neural mode (**Fig. 3c**). Both *pcdh19^-^*^10b^*^p^* and *pcdh19^Δprom^* showed decreased correlation with the group average (**Fig. 3c**), although this varied by neural mode between the two mutants. The *pcdh19^-^*^10b^*^p^* larvae showed the greatest change along the second neural mode, while *pcdh19^Δprom^* mutants exhibited significant differences along neural modes 2-5 (**Fig. 3c**). The response to drifting sinusoidal gratings diverged from wild type for both *pcdh19* mutants, as the trajectories of individual mutant fish were less correlated to the wild type average than were individual wild type fish. To determine whether individual mutants described similar or distinct trajectories, we calculated pairwise correlations between larvae within an experimental group. If mutants converged on a distinct, but stereotyped, circuit organization, then their intra-group correlation should be similar to the intra-group correlations of the wild type group. Alternatively, if the divergence of the mutants from wild type is due to stochastic variations, then the intra-group correlations should be lower and exhibit an increased variance. Consistent with stochastic variation, the *pcdh19* mutants exhibit lower intra-group correlations than wild type larvae (**Fig. 3d**). When visualized in three dimensions, the trajectories from individual *pcdh19^Δprom^* mutants differ dramatically from the stereotyped paths of wild type larvae (**Fig. 2g,h**), and diverge from each other (**Fig. 3e,f**). However, despite the variability of the individual mutants, averaging within a genotype suppresses this variation and gives rise to averaged dynamics that are strongly correlated with wild type (**Fig. 3g**). This suggests that the effect of *pcdh19* loss on visual processing is best understood as stochastic developmental noise, as this noise can be suppressed by averaging (**Fig. 3g**).

**Figure 3.**
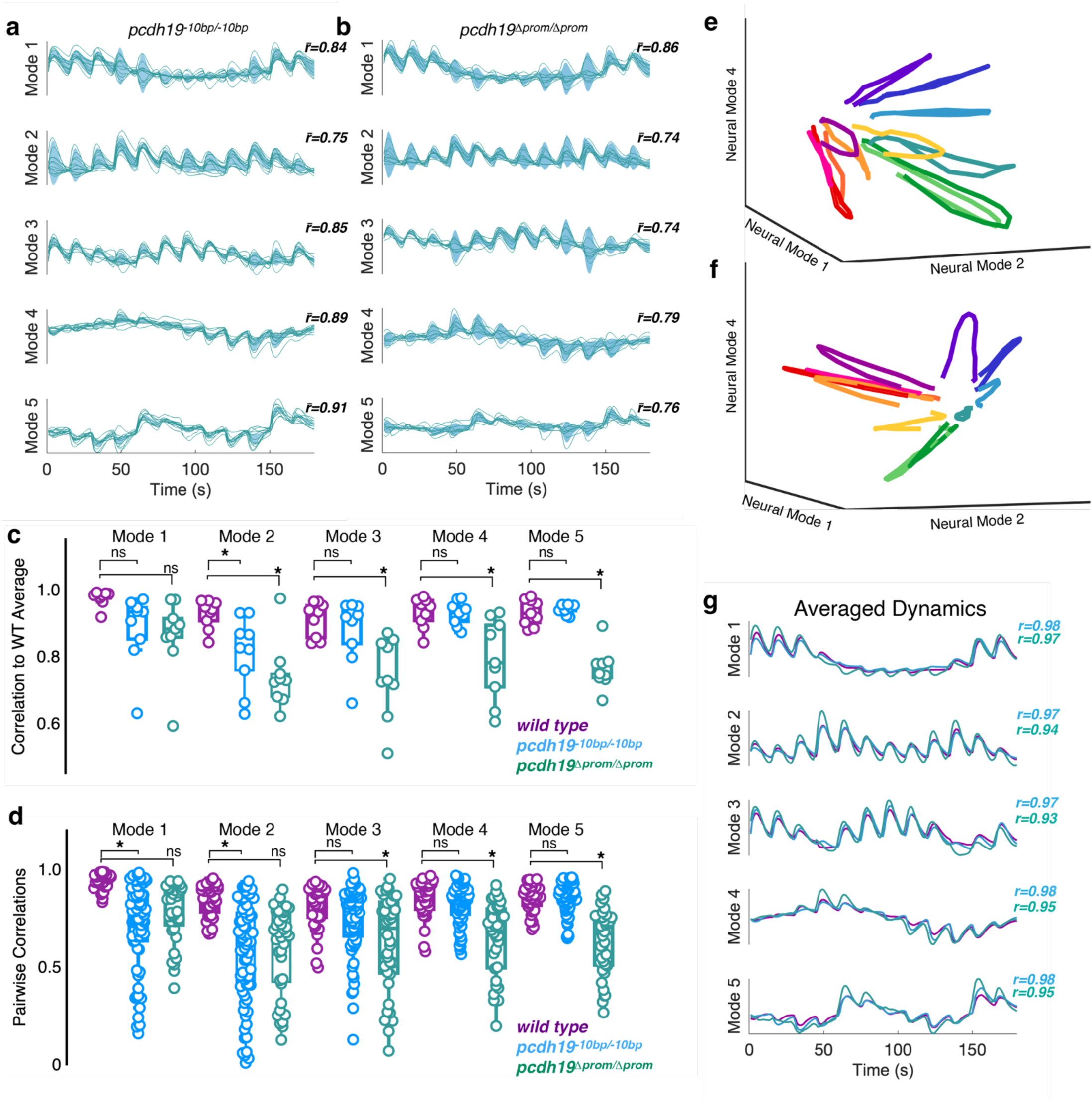
Mutations in pcdh19 lead to altered neural dynamics. **a,b.** Shown are latent dynamics determined from trial averaged neural data collected in homozygous pcdh19^-^^10b^^p^ (**a**) and pcdh19^Δprom^ (**b**) mutant larvae. Individual traces are in teal and the shaded area in blue indicates the group variance. The value, *r*’, represents the mean correlation of each individual neural mode to the group averaged neural mode. We obtained a total of 3589 visually responsive neurons from pcdh19^-^^10b^^p^ larvae, and 2172 neurons from pcdh19^Δprom^ larvae. **c.** Boxplot shows the correlation of individual trial-averaged neural modes to the wild type average (wild type, n=9: 0.97±0.008, mode 1; 0.93±0.014, mode 2; 0.91±0.017, mode 3; 0.93±0.015, mode 4; 0.93±0.013, mode 5; pcdh19^-^^10b^^p^, n=12: 0.84±0.035, mode 1; 0.75±0.03, mode 2; 0.85±0.027, mode 3; 0.89±0.014, mode 4; 0.91±0.015, mode 5; pcdh19^Δprom^, n=9: 0.88±0.03, mode 1; 0.78±0.04, mode 2; 0.77±0.046, mode 3; 0.80±0.04, mode 4; 0.78±0.022, mode 5; *p<0.05, Dunnett’s test). **d.** Pairwise within-group correlations of neural modes for each genotype (wild type,n=36: 0.94±0.013, mode 1; 0.84±0.028, mode 2; 0.81±0.038, mode 3; 0.85±0.031, mode 4; 0.85±0.023, mode 5; pcdh19^-^ ^10b^^p^, n=66: 0.71±0.064, mode 1; 0.56±0.076, mode 2; 074±0.051, mode 3; 0.82±0.03, mode 4; 0.86±0.022, mode 5 ; pcdh19^Δprom^, n=36: 0.78±0.05, mode 1; 0.59±0.075, mode 2; 0.61±0.095, mode 3; 0.64±0.031, mode 4; 0.63±0.056, mode 5; *p<0.05, Dunnett’s test). **e,f.** Three-dimensional representations of latent dynamics of individual pcdh19^Δprom^ mutant larvae. The view is the same as in Fig. 2g**-i**. **g.** Neural modes were averaged within each genotype (wild type/purple, pcdh19^-^^10b^^p^/blue, pcdh19^Δprom^/teal. The value, *r*, represents the correlation of each mutant average with the wild type average (pcdh19^-^^10b^^p^/blue, pcdh19^Δprom^/teal).

Pcdh19 is a member of a small family of related δ2-pcdhs. To determine if mutations in other δ2-pcdhs also affect visual responses in the optic tectum, we recorded the response to visual stimulation in mutants lacking *pcdh17* (*pcdh17^-^*^5b^*^p^*)^37^. As above, we used the trial averaged neuronal responses to compute the latent dynamics along the first five neural modes (**Fig. 4a**). Similar to the *pcdh19* mutants, the latent dynamics exhibited by *pcdh17^-^*^5b^*^p^* mutants was less correlated with the wild type average (**Fig. 4a**), although the differences were not statistically significant (**Fig. 4b**). To measure the intra-group variation exhibited by the *pcdh17^-^*^5b^*^p^* mutants, we determined pairwise correlations along each neural mode (**Fig. 4c**). Although there was a uniform decrease in pairwise correlation among the *pcdh17^-^*^5b^*^p^* mutants, this was only significant for neural mode 5 (**Fig. 4c**). This observed intra-group noise was eliminated by averaging, as the *pcdh17^-^*^5b^*^p^* group average exhibited high correlation with the wild type average (**Fig 4d**). As each mutant genotype showed a reduced intra-group correlation, it is possible that this increased variation is due to unreliable or noisy neuronal responses. To determine whether the tectal responses to visual stimulation are reliable, we determined the between-trial correlations for each neural mode within each experimental group (**Fig. 4e**). We found no significant difference in the between-trial consistency of the population dynamics in the mutant groups, compared to wild type (**Fig. 4e**). This suggests that each mutant network is responding reliably and reproducibly to repeated presentations of a given stimulus, yet deviates randomly from the wild type. To quantify the within-group variance, we generated histograms of all pairwise correlation coefficients within each experimental group (**Fig. 4f**). While the distribution of correlation coefficients for the wild type group is narrow and concentrated at higher correlations, the mutants all exhibit significantly broader distributions (**Fig. 4f**). To provide a metric for the overall similarity of latent dynamics among different larvae, we calculated the mean *r* of individual correlations across the five neural modes for each genotype (**Fig. 4g**). If the computed dynamics are empirical estimates of the neural manifold, then this summary *r* should measure the similarity of an individual manifold to the averaged wild type manifold. In the case of each mutant genotype, the dynamics are significantly different from wild type, although the effect is more robust for the *pcdh19^Δprom^* mutants (**Fig. 4g**). While the variations along individual neural modes are not considered to be statistically significant for the *pcdh17^-^*^5b^*^p^* mutants, the aggregate impact on the shape of the neural manifold is significant (**Fig. 4g**). Thus, loss of either *pcdh19* or *pcdh17* alters the neural population response to visual stimulation, by introducing stochastic perturbations into circuit assembly. Our data are consistent with the idea that δ2-pcdhs contribute to canalization of the developing optic tectum (**Fig. 4h**). To determine the relationship of the altered latent dynamics in the pcdh mutants to more traditional measures of neuronal responses, we determined the orientation and direction selectivity for neurons in wild type larvae and each of the mutants (**Fig. S2**). For each neuron, we determined an orientation selectivity index (OSI), direction selectivity index (DSI) and preferred direction. The the *pcdh17^-^*^5b^*^p^* mutants exhibited a dramatic decrease in orientation selectivity (**Fig. S2a**) and direction selectivity (**Fig. S2b**), while both *pcdh19^-^*^10b^*^p^* and *pcdh19^Δprom^* showed a shift toward increased orientation selectivity and a modest decrease in direction selectivity in the *pcdh19^Δprom^* mutants. The distribution of preferred orientations also exhibited differences among the mutants, as *pcdh17^-^*^5b^*^p^* mutants exhibited a decreased response to rostral-to-caudal motion (**Fig. S2c,d**), and both *pcdh19^-^*^10b^*^p^* (**Fig. S2c,e**) and *pcdh19^Δprom^* (**Fig. S3c,f**) showed an increase in the proportion of neurons responding to rostral-to-caudal motion.

**Figure 4.**
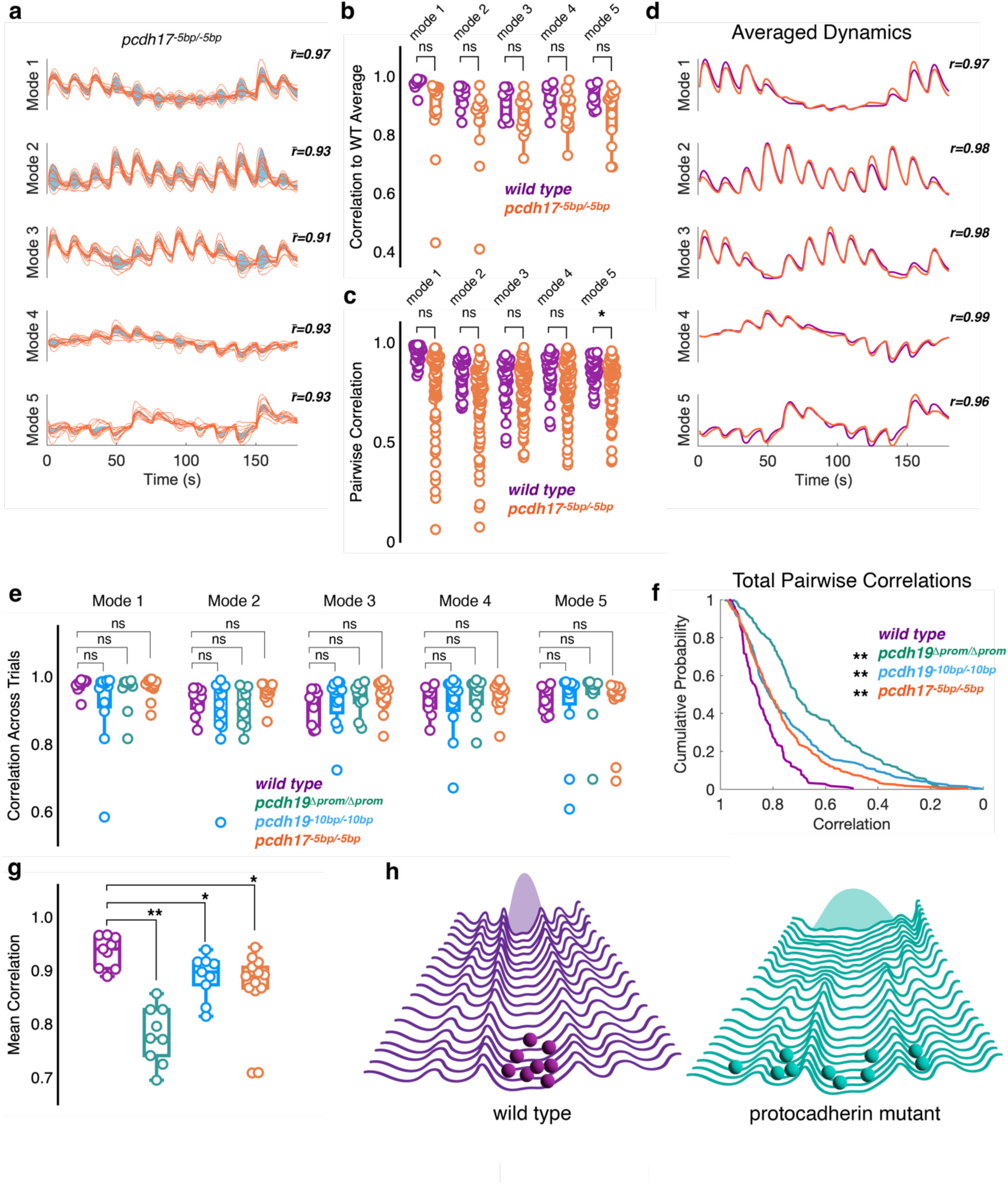
Mutations in pcdh17 lead to altered neural dynamics. **a.** Latent dynamics computed from trial averaged neural data collected in homozygous pcdh17^-5b^^p^ mutant larvae. Individual traces are in orange and the shaded area in blue indicates the group variance. The value, *r*’, represents the mean correlation of each individual neural mode to the group average of that neural mode. We obtained a total of 3561 visually-responsive neurons. **b.** The correlation of individual trial-averaged neural modes to the wild type average (wild type: 0.97±0.008, mode 1; 0.93±0.014, mode 2; 0.91±0.017, mode 3; 0.93±0.015, mode 4; 0.93±0.013, mode 5, n=9; pcdh17^-5b^^p^: 0.87±0.042, mode 1; 0.83±0.04, mode 2; 0.87±0.019, mode 3; 0.89±0.027, mode 4; 0.87±0.093, mode 5, n=13). **c.** Pairwise within-group correlations of neural modes for wild type (purple) and pcdh17 mutants (orange). (wild type, n=36: 0.94±0.013, mode 1; 0.84±0.028, mode 2; 0.81±0.038, mode 3; 0.85±0.031, mode 4; 0.85±0.023, mode 5; pcdh17^-5b^^p^, n=78: 0.78±0.055, mode 1; 0.71±0.053, mode 2; 0.77±0.037, mode 3; 0.79±0.039, mode 4; 0.79±0.035, mode 5; *p,0.05, Dunnett’s test). **d.** Neural modes were averaged for pcdh17^-5b^^p^ mutants (orange) and the average was compared to the wild type average (purple). The value, *r*’, represents the correlation of each mutant average with the wild type average. **e.** The mean trial to trial correlations are shown for wild type (purple), pcdh19^-^^10b^^p^ (blue) pcdh19^Δprom^ (teal), and pcdh17^-5b^^p^ (orange) mutants. Trial-to-trial variability in the mutants is not statistically different from wild type larvae (wild type, n=9: 0.98±0.006, mode 1; 0.94±0.012, mode 2; 0.96±0.01, mode 3; 0.95±0.017, mode 4; 0.94±0.032, mode 5; pcdh19^-^^10b^^p^: 0.92±0.034, mode 1; 0.89±0.034, mode 2; 0.92±0.022, mode 3; 0.92±0.027, mode 4; 0.91±0.036, mode 5; pcdh19^Δprom^: 0.91±0.024, mode 1; 0.90±0.025, mode 2; 0.90±0.025, mode 3; 0.90±0.015, mode 4; 0.85±0.049, mode 5; pcdh17^-5b^^p^, n=13: 0.97±0.009, mode 1; 0.95±0.009, mode 2; 0.94±0.013, mode 3; 0.94±0.012, mode 4; 0.92±0.026, mode 5). **f.** Cumulative probability histogram for the distribution of all pairwise within group *r* values for wild type (purple), pcdh19^-^^10b^^p^ (blue) pcdh19^Δprom^ (teal), and pcdh17^-5b^^p^ (orange) mutants. (**p<0.0001, Dunn’s test). **g.** Mean correlation (*r*) of individual datasets to the wild type average, averaged across neural modes (wild type, n=9: 0.90±0.018; pcdh19^-^^10b^^p^, n=12: 0.83±0.017; pcdh19^Δprom^, n=9: 0.74±0.033; pcdh17^-5b^^p^, n=13: 0.82±0.028;*p<0.05, **p<0.0001, Dunnett’s test). **h.** Hypothesized roles of Pcdh19 and Pcdh17 in canalyzing the assembly of visual circuits in the zebrafish optic tectum. In the wild type scenario (left/purple), development is canalized and the outcomes (circuit response to visual stimulation) is restricted, showing a small distribution in outcomes (shaded Gaussian). In the case of protocadherin mutants (right/teal), the developmental landscape is flattened, allowing development to stochastically explore a variety of alternative paths, leading to an increased variation among outcomes (broad shaded distribution).

In addition to homozygous mutants, we also analyzed the visual responses of heterozygous mutants (**Fig. S3**). Latent dynamics in the heterozygotes was similar to each of the respective homozygous mutants (**Fig. S3a-c**). Both the correlation of individual neural modes to the wild type average (**Fig. S3d**) and the pairwise, within-group correlations (**Fig. S3e**) were lower than for wild type and the reduction was comparable in magnitude to what we observed in homozygous mutants. When averaged across neural modes, the mean similarity of each δ2-pcdh mutant to the wild type neural response was reduced (**Fig. S3f**), and was characterized by increased within-group variation (**Fig. S3g**). As was observed for homozygous mutants (**Fig. 3,4**), this variability could be averaged away and averages of the heterozygous mutants were highly correlated with the wild type average (**Fig. S3h,i**).

## Discussion

The δ2-pcdhs comprise a subfamily of homophilic cell adhesion molecules within the cadherin superfamily, and are associated with neurodevelopmental disorders in humans^40,41^. Among the five δ2-pcdhs in mammals, all are linked to human disorders: *PCDH8* and *PCDH10* are linked to autism^42,43^, a genomic deletion encompassing *PCDH18* caused severe intellectual disability^44^, *PCDH17* is implicated in mood disorders^45^, and mutations in *PCDH19* cause a developmental epileptic encephalopathy^22,23^, and are associated with autism and schizophrenia ^20,21^. We previously used large-scale *in vivo* calcium imaging of spontaneous activity and network analysis to show that zebrafish mutants lacking *pcdh19* had defects in network topology^46^. Subsequently, others showed that mutant rodents and zebrafish larvae were hyperexcitable^47,48^, and that mutant rodents showed increased network synchrony^47^. Here, our results show that loss of either *pcdh19* or *pcdh17* impairs visual processing in the optic tectum of the larval zebrafish.

Our data show that the response of the larval optic tectum to moving sinusoidal gratings is encoded by low-dimensional dynamics that are confined to a neural manifold. These dynamics are highly stereotyped among wild type individuals. It was recently shown that dynamics are preserved across individuals within a species, but these dynamics need to be aligned using a procedure such as canonical correlation analysis^34^, due to the fact that the latent dynamics are estimated from only a small subsample of neurons that vary among animals. Here, we image a substantial proportion of the visually responsive neurons in the optic tectum. Consequently, the dynamics derived from different individuals do not require further alignment. This allows the complex neural responses of hundreds of neurons in different animals to be reduced to a common set of activity patterns that can be quantitatively compared within and between genotypes. The variation among wild type individual larvae is very low, indicating that development within a population gives rise to an ensemble of equivalent networks that differ in the precise number, morphology and connectivity of the constituent neurons, but whose overall functional variation is low.

Elimination of either *pcdh19* or *pcdh17* leads to altered latent dynamics. The stereotyped neural response that characterizes the population of wild type larvae is degraded in the mutants, whose dynamics are less well correlated to the averaged wild type response. The latent dynamics among the mutant genotypes varies by individual. As a consequence, within a mutant group, individuals are much less similar to each other than the pool of wild type larvae are to each other. Additionally, this stochastic variability is masked in a group average, as the deviations specific to an individual can be averaged away. This implies that the loss of Pcdh19 or Pcdh17 does not lead to a particular, alternative network structure, but to a stochastic array of alternatives. Thus, the effects of the *pcdh19* and *pcdh17* mutations are best understood as an increase in developmental noise, leading to an increase in phenotypic variability. Our data show that the effects of a well-defined genetic lesion on a simple, highly stereotyped network can be stochastic. While both *pcdh19^-^*^10b^*^p^* and *pcdh19^Δprom^* mutants result in a complete loss of Pcdh19 protein, their effects on the optic tectum are not identical. The phenotype associated with Pcdh19 loss in the *pcdh19^Δprom^* line is significantly more severe than in the *pcdh19^-^*^10b^*^p^* line. Combined with the observation that the magnitude of the impact of indel mutations in *pcdh19^-^*^10b^*^p^* and *pcdh17^-^*^5b^*^p^* is similar, this suggests that the difference could be due to the activation of compensatory mechanisms by the indel mutations that are absent in the promoterless allele. This may indicate a further source of variability in understanding neural disorders and in the study of relevant genes in animal models, as even alternative “null” mutations may give rise to distinct phenotypes.

In evolution, strong selective pressures prevent the accumulation of mutations that have an adverse impact on phenotypic fitness. Release of a selective pressure can then lead to an increase in allelic and phenotypic diversity within a population. By analogy, our results suggest that Pcdh19 and Pcdh17 suppress variation in the development of tectal circuitry. Loss of either protocadherin releases a constraint on circuit assembly, allowing stochastic variation in circuit organization and function. These results suggest that Pcdh19 and Pcdh17 contribute to developmental robustness by canalizing circuit assembly and limiting developmental variability. While canalization is most commonly discussed in terms of gene regulatory networks buffering against adverse genetic or environmental effects^49,50^, we suggest that the protocadherins may buffer against the formation of inappropriate connections and help guide circuit assembly toward an optimal outcome. These results suggest that while the heterogeneity observed in neurodevelopmental disorders is often attributed to polygenic effects, it could also be due to decanalization of circuit assembly and stochastic degradation of circuit function.

## Methods

### Zebrafish maintenance and generation of lines

Adult zebrafish (Danio rerio) were maintained at ∼28.5°C and staged according to Westerfield (1995). All animal procedures were performed in accordance with the Ohio State University animal care committee’s regulations. We previously generated lines harboring germline mutations of *pcdh19* (*pcdh19^os^*^51^)^36^ and *pcdh17* (*pcdh17^os^*^69^)^37^.

The *pcdh19^Δprom^* mutant line (*pcdh19^os^*^77^) was established by co-injecting ribonucleoprotein complexes (Integrated DNA Technologies) consisting of Cas9 protein and gRNAs targeting a site in the 5’ end of *pcdh19* exon1 (GGGCTCAGATTAACCCATCG) and ∼4 kb upstream of the ATG start codon (CTGTTGTGAGCTAGTTACCA), which is predicted to eliminate the promoter region. Founders were identified by PCR and sequenced, confirming the large genomic deletion. Like the *pcdh19^-^*^10b^*^p^* line, Western blots confirmed that the homozygous *pcdh19^Δprom^* line produced no Pcdh19 protein.

The plasmid *Tol2-elavl3:H2B-GCaMP8s* was assembled using the following plasmids: Tol2-elavl3-GCaMP6s (a gift from Misha Ahrens; Addgene plasmid # 59530 ; http://n2t.net/addgene:59530 ; RRID:Addgene_59530), CMV:Histone H2B-mGL (H2B-mGreenLantern was a gift from Gregory Petsko; Addgene plasmid # 164464 ; http://n2t.net/addgene:164464 ; RRID:Addgene_164464), and CMV:jGCaMP8s (gift from GENIE Project; Addgene plasmid # 162371 ; http://n2t.net/addgene:162371 ; RRID:Addgene_162371). Along with mRNA encoding Tol2 transpose, this plasmid was co-injected into 1 cell stage *nacre* embryos in order to establish the transgenic line *Tg(elavl3:H2B-GCaMP)^os^*^78^. This transgenic line was crossed with the *pcdh19^-^*^10b^*^p^*, *pcdh19^Δprom^*and *pcdh17^-^*^5b^*^p^* mutant lines.

### Calcium imaging

Unanaesthetized 6 days-post-fertilization (dpf) larvae were embedded dorsal side up in 2% low melting point agarose. Imaging was performed on a custom-built resonant-scanning 2-photon microscope. Briefly, 920nm excitation was provided by an Axon920-TPC laser (Coherent, Inc.). We used a Nikon Apochromat 25x/1.1NA water-immersion objective for imaging. The resonant scan-head and controller, 3DMS robotic stage, Hamamatsu GaAsP photomultiplier tubes and power supply were obtained from Sutter Instruments (www.sutter.com). A piezo-electric objective positioner (nPFocus250) was obtained from nPoint (www.npoint.com). The microscope was run with ScanImage 5.2^51^ (Vidrio Technologies). All other parts were obtained from Thorlabs (www.thorlabs.com). Image stacks of 7 optical sections (512×512), spaced at 10 μm, were collected at 1 s intervals, with a pixel size of 0.65 μm. For each group, we included data from 9-13 larvae, which is in line with the number of fish used in comparable studies^52–54^. These were derived from at least two, separate crosses.

### Visual stimulation

Visual stimuli were programmed in Python, using PsychoPy3 (https://psychopy.org). Moving sinusoidal gratings were presented for 5 seconds, with 10 seconds of a neutral gray background in between each direction, which were rotated at 30° intervals. The stimulus set was presented three times, with 50 seconds in between each presentation. Stimuli were projected onto a translucent screen 2.5 cm from the larvae using a Rif6 cube picoprojector (rif6.com). The stimulus occupied ∼104° of the visual field.

### Data analysis

We used CaImAn-MATLAB for processing of calcium imaging movies^28^, including non-rigid motion correction with NoRMCorre^55^. First, sequences of image stacks (735 stacks of 7 planes) were re-formatted to movies of image planes (7 stacks of 735 timepoints). These were motion corrected and inspected for motion artifacts. Movies that exhibited *z*-drift or other defects were discarded. Brightness and contrast were adjusted and then images were masked in order to analyze only the left (visually-responsive) hemi-tectum. CaImAn was used to segment movies and extract ΔF/F fluorescence traces. Cells that were responsive to the visual stimuli in each of the three trials were selected for further analysis.

Principal Component Analysis was performed on individual datasets. As ∼90% of the variance was explained by the top 5 principal components (neural modes), we restricted our analysis to these. We projected the original neural responses onto these new basis vectors. The order of neural modes varied among datasets, as did the sign of the neural mode. We established a standard order and inspected each dataset to ensure a common order and sign for the neural modes. For comparisons of neural modes among datasets, we used Pearson’s correlation, *r*^34,35^. First, we averaged dynamics along each neural mode to generate a wild type average. Correlation of neural modes from individual larvae to the averaged neural modes provided a measure of variation. As an alternative measure of the variation in the datasets, each individual set of neural modes was correlated to all others within a group. This pairwise correlation describes how much latent dynamics varied among individuals within a genotype. To generate a single measure of similarity of latent dynamics between an individual and the wild type average, we averaged the correlation coefficients across the 5 neural modes. For comparisons, statistical significance was determined using Dunnett’s test. For the comparison of distributions of *r*, we used a Dunn’s test. To compare circular distributions, we used a MATLAB implementation of Watson’s *U*^2^ test (Pierre Mégevand (2025) pierremegevand/watsons_u2 https://github.com/pierremegevand/watsons_u2). All other statistical analysis was performed in JMP Pro 17.

### Canonical Correlation Analysis

CCA was implemented in MATLAB and performed as described in Gallego et al. (2018). Briefly, a QR decomposition was performed on each set of dynamics to be aligned (*L*_1_ = *Q*_1_*R*_1_ and *L*_1_ = *Q*_2_*R*_2_), then a singular value decomposition was performed on the inner product of *Q*_1_ and 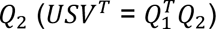. The dynamics were then aligned by 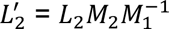, where 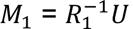 and 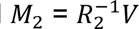.

## Acknowledgements

The content is solely the responsibility of the authors and does not necessarily represent the official views of the National Institutes of Health. Research reported in this publication was supported by the National Eye Institute of the National Institutes of Health under award number R21EY034706 and by the National Insitute of General Medical Sciences of the National Institutes of Health under award R01GM141280 to JDJ. We would like to thank Marcus Nichols for support with zebrafish maintenance and husbandry.

---

**Supplemental Figure 1.**
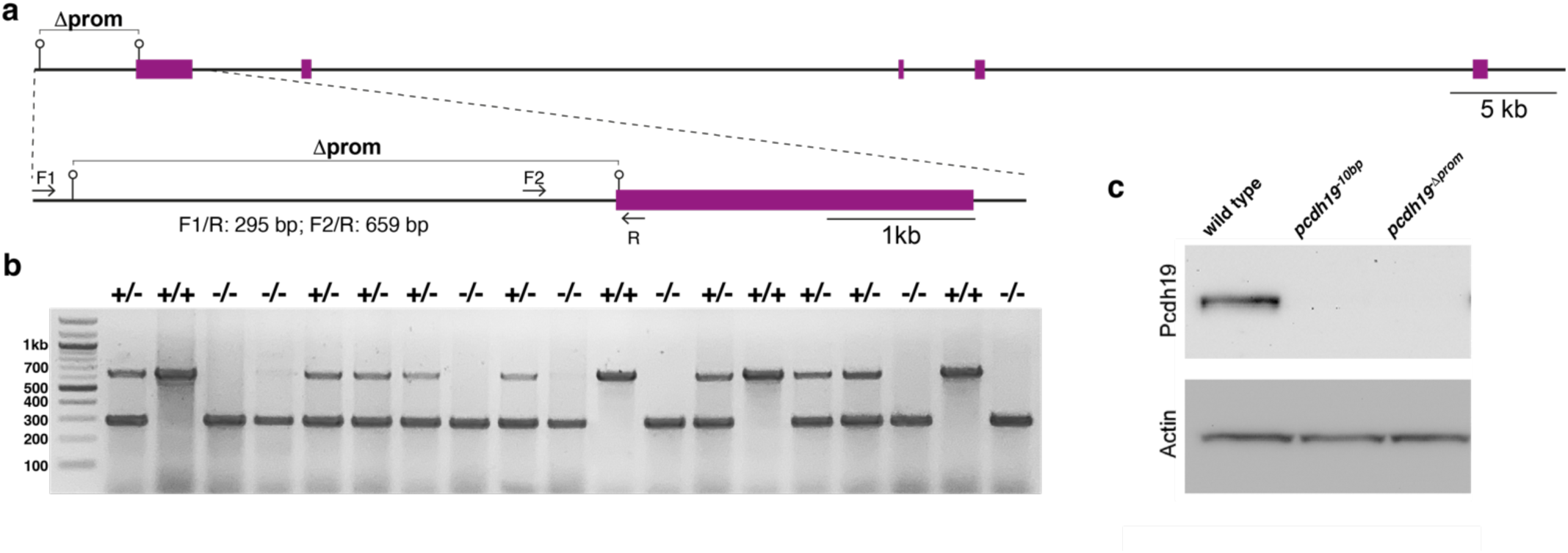
**a** Schematic of the zebrafish pcdh19 gene, with exons shown in purple. Pins show the sites of CRISPR/Cas9 target sites. Primers used for PCR-based screening are shown, as are the expected sizes of the PCR products. **b** Genotyping of mutant in-crosses using F1, F2 and R primers, which distinguish wild type, heterozygous and homozygous mutant larvae. **C** Western blot showing the absence of Pcdh19 protein in both our previously published mutant line and in the promoterless line presented here.

**Supplemental Figure 2.**
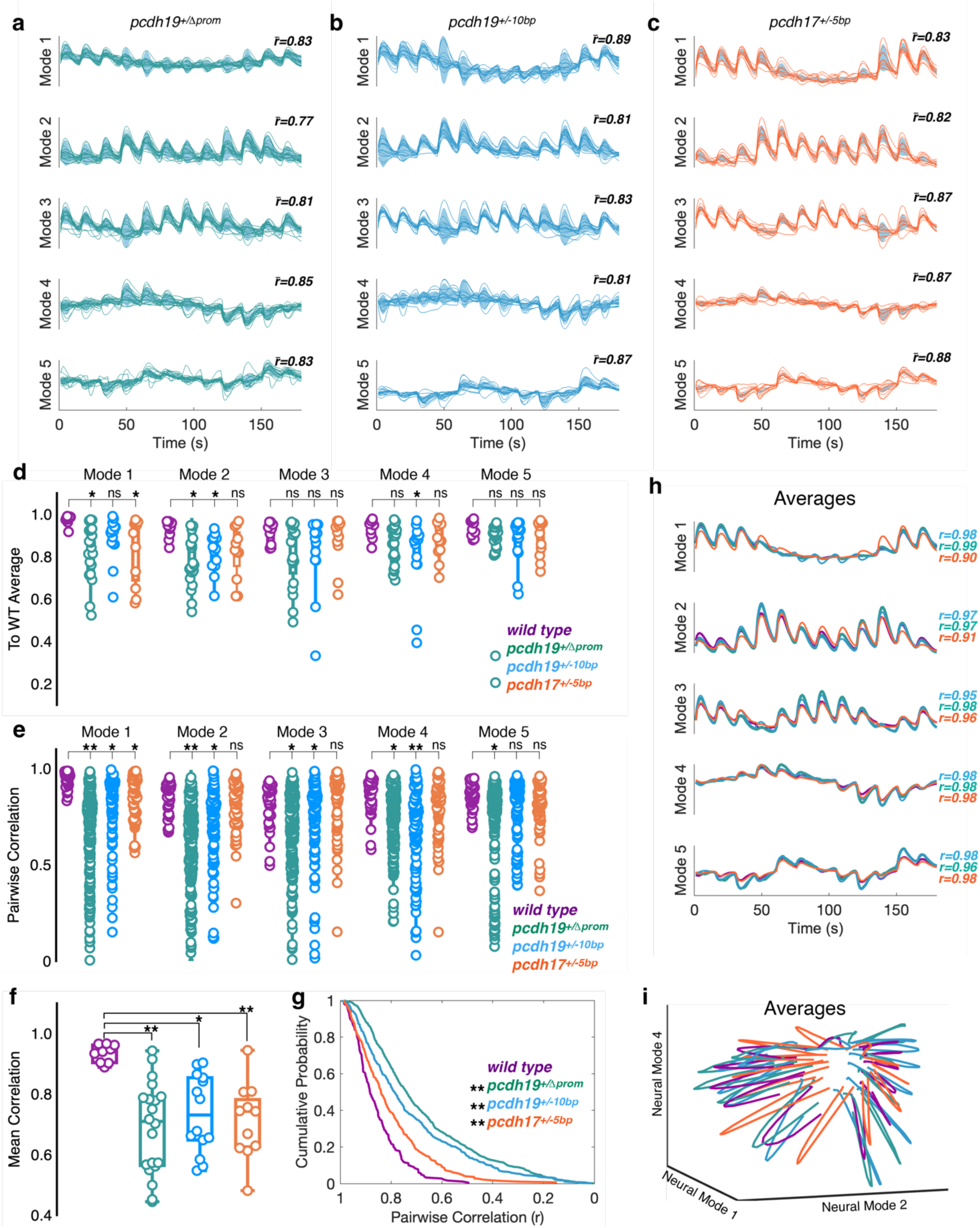
Heterozygous δ2-pcdh mutants exhibit altered neural dynamics. **a-c.** Latent dynamics computed from trial averaged neural data collected in heterozygous (**a**) pcdh19^Δprom^, (**b**) pcdh19^-^^10b^^p^ and (**c**) pcdh17^-5b^^p^ mutant larvae. Individual traces are shown on top of the group variance, shaded in blue. The value, *r*’, represents the mean correlation of each individual neural mode to the wild type average of that neural mode. **d.** The correlation of individual trial-averaged neural modes to the wild type average (wild type: 0.97±0.008, mode 1; 0.93±0.014, mode 2; 0.91±0.017, mode 3; 0.93±0.015, mode 4; 0.93±0.013, mode 5, n=9; pcdh19^Δprom^: 0.82±0.029, mode 1; 0.77±0.026, mode 2; 0.81±0.033, mode 3; 0.85±0.02, mode 4; 0.83±0.044, mode 5, n=20; pcdh19^-^^10b^^p^ : 0.89±0.027, mode 1; 0.81±0.24, mode 2; 0.83±0.047, mode 3; 0.81±0.047, mode 4; 0.87±0.029, mode 5, n=14; pcdh17^-5b^^p^: 0.83±0.048, mode 1; 0.82±0.038, mode 2; 0.87±0.036, mode 3; 0.87±0.027, mode 4; 0.88±0.026, mode 5, n=11). (*p<0.05, Dunnett’s test). **e.** Pairwise within-group correlations of neural modes for wild type (purple) and pcdh17 mutants (orange). (wild type, n=36: 0.94±0.007, mode 1; 0.84±0.014, mode 2; 0.81±0.019, mode 3; 0.85±0.016, mode 4; 0.85±0.011, mode 5; pcdh19^Δprom^, n=190: 0.66±0.017, mode 1; 0.60±0.016, mode 2; 0.66±0.015, mode 3; 0.74±0.012, mode 4; 0.73±0.016, mode 5 pcdh19^-^^10b^^p^, n=91: 0.8±0.02, mode 1; 0.68±0.022, mode 2; 0.72±0.024, mode 3; 0.65±0.026, mode 4; 0.78±0.017, mode 5; pcdh17^-5b^^p^, n=55: 0.84±0.016, mode 1; 080±0.017, mode 2; 0.81±0.0223, mode 3; 0.79±0.022, mode 4; 0.79±0.018, mode 5; **p<0.0001, *p,0.05, Dunnett’s test). **f.** Neural modes were averaged for wild type (purple, 0.9±0.018), pcdh19^-^^10b^^p^ (blue, 0.74±0.034), pcdh19^Δprom^ (teal, 0.69±0.041) and pcdh17^-5b^^p^ mutants (orange, 0.71±0.038) and the average was compared to the wild type average (purple). (**p<0.0001, *p<0.05, Dunnett’s test). **g.** Cumulative probability histogram for the distribution of all pairwise within group *r* values for wild type (purple), pcdh19^-^^10b^^p^ (blue) pcdh19^Δprom^ (teal), and pcdh17^-5b^^p^ (orange) mutants. (**p<0.0001, Dunn’s test). **h.** Mean correlation (*r*) of individual datasets to the wild type average, averaged across neural modes (wild type, n=9: 0.90±0.018; pcdh19^-^^10b^^p^, n=12: 0.83±0.017; pcdh19^Δprom^, n=9: 0.74±0.033; pcdh17^-5b^^p^, n=13: 0.82±0.028;*p<0.05, **p<0.0001, Dunnett’s test). **i.** Plotting of averaged dynamics along neural modes 1, 2 and 4 for wild type (purple), pcdh19^-^^10b^^p^ (blue) pcdh19^Δprom^ (teal), and pcdh17^-5b^^p^ (orange) heterozygous mutants.

**Supplemental Figure 3.**
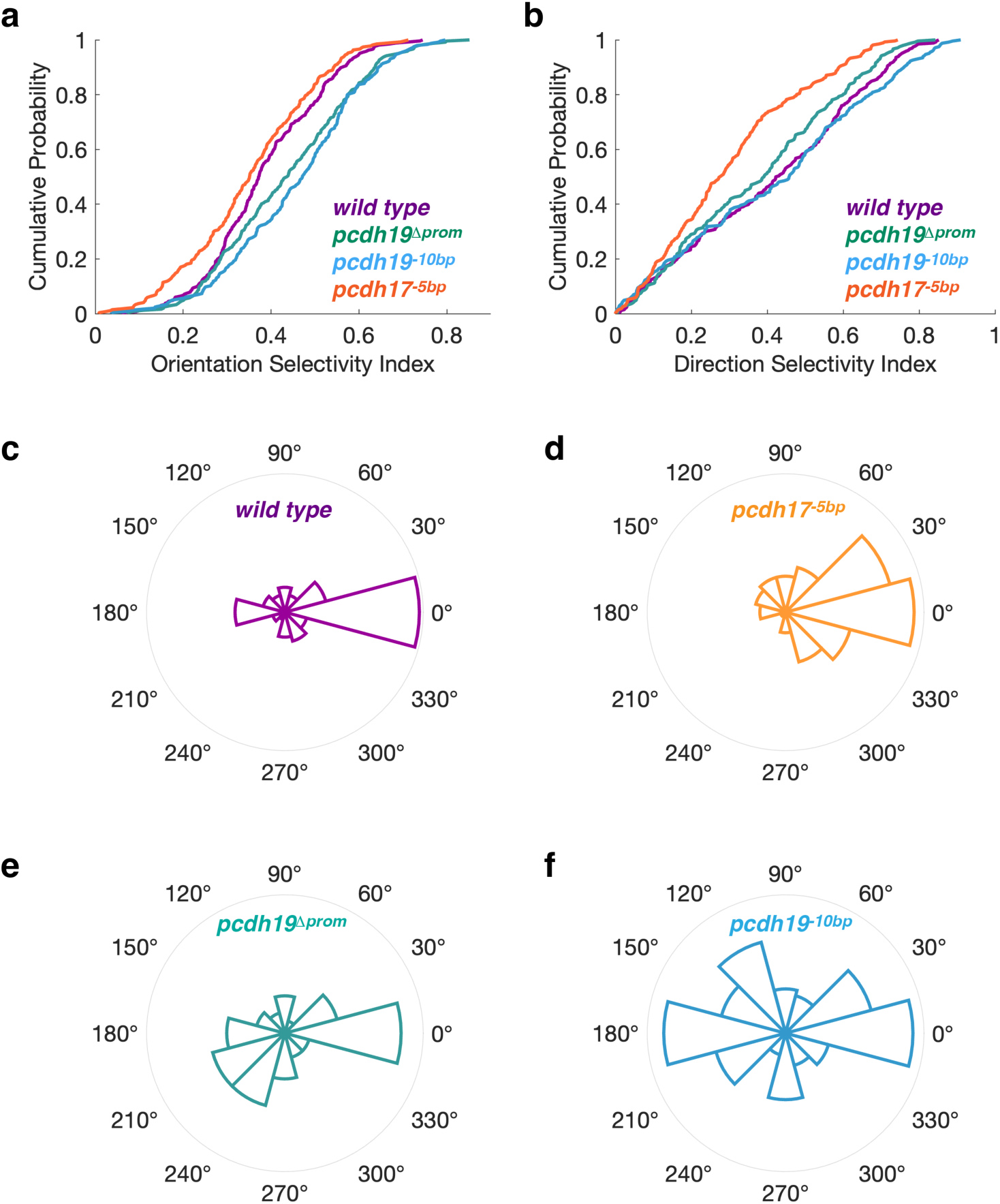
Altered orientation selectivity and direction selectivity in δ2-pcdh mutants. **a.** Cumulative probability distributions of orientation selectivity for all neurons in wild type (purple), pcdh19^-^^10b^^p^ (blue) pcdh19^Δprom^ (teal), and pcdh17^-5b^^p^ (orange) mutants. Pcdh17 and Pcdh19 mutants exhibit distinct effects on orientation selectivity, with pcdh17^-5b^^p^ mutants exhibiting an overall shift toward reduced orientation selectivity, and both pcdh19^-^^10b^^p^ and pcdh19^Δprom^ showing a shift toward increased direction selectivity.(**p<0.0001; Dunn’s test). **b.** Cumulative probability distributions of direction selectivity for all neurons in wild type (purple), pcdh19^-^^10b^^p^ (blue) pcdh19^Δprom^ (teal), and pcdh17^-5b^^p^ (orange) mutants. The pcdh17^-5b^^p^ mutants exhibit a dramatic shift toward reduced direction selectivity. In contrastboth pcdh19^-^^10b^^p^ and pcdh19^Δprom^ showing a shift toward increased direction selectivity.(*p<0.05, **p<0.0001; Dunn’s test). **d.** A polar histogram showing the preferred directions of direction selective neurons (DSI>0.33) in wild type larvae. The preferred direction as obtained as the vector average of the responses across all stimuli. **e.** A polar histogram showing the preferred direction of direction selective neurons in pcdh17^-5b^^p^ mutants. There is an increase in the number of cells that prefer caudal-to-rostral motion (p<0.0001; Watson’s U2 test). **f.** A polar histogram showing the preferred direction of direction selective neurons in pcdh19^Δprom^ mutants. There is a slight decrease in cells responsive to caudal-to-rostral motion and an increase in the number of cells that prefer motion between 195°-240° (p<0.0001; Watson’s U2 test). **g.** A polar histogram showing the preferred direction of direction selective neurons in pcdh19^-^^10b^^p^ mutants. There is a large increase in the proportion of cells responsive to directions other than caudal-to-rostral motion, particularly directions centered around 120° and 180° (p<0.0001; Watson’s U2 test).

